# Functional differences in the hypothalamic-pituitary-gonadal axis are associated with alternative reproductive tactics based on an inversion polymorphism

**DOI:** 10.1101/2020.09.08.285676

**Authors:** JL Loveland, LM Giraldo-Deck, D Lank, W Goymann, M Gahr, C Küpper

**Author notes:** Corresponding author: Jasmine L. Loveland.

## Abstract

The evolution of social behavior depends on genetic changes, yet, how genomic variation manifests itself in behavioral diversity is still largely unresolved. Chromosomal inversions can play a pivotal role in producing distinct behavioral phenotypes, in particular, when inversion genes are functionally associated with hormone synthesis and signaling. Male Ruffs exhibit alternative reproductive tactics (ARTs) with an autosomal inversion determining two alternative morphs with clear behavioral and hormonal differences to the ancestral morph. We investigated hormonal and transcriptomic differences in the pituitary and gonads. Using a GnRH challenge, we found that the ability to synthesize testosterone in inversion carriers is severely constrained, whereas the synthesis of androstenedione, a testosterone precursor, is not. Inversion morphs were able to produce a transient increase in androstenedione following the GnRH injection, supporting the view that pituitary sensitivity to GnRH is comparable to that of the ancestral morph. We then performed gene expression analyses in a second set of untreated birds and found no evidence of alterations to pituitary sensitivity, gonadotropin production or gonad sensitivity to luteinizing hormone or follicle-stimulating hormone across morphs. Inversion morphs also showed reduced progesterone receptor expression in the pituitary. Strikingly, in the gonads, inversion morphs over-expressed *STAR*, a gene that is located outside of the inversion and responsible for providing the cholesterol substrate required for the synthesis of sex hormones. In conclusion, our results suggest that the gonads determine morph-specific differences in hormonal regulation.

## Introduction

Genomic rearrangements such as chromosomal inversions frequently provide the genetic variation necessary for the evolution of behavioral diversity associated with mating and reproduction (Chouteau et al., 2017; Gilburn and Day, 1999; Horton et al., 2014b; Küpper et al., 2016; Pearse et al., 2014). At the proximate level, these so-called “supergenes” may contribute to hormonal plasticity by capturing genes required for hormone production and sensitivity. Over time, sequence evolution combined with selection on certain inversion haplotypes may further canalize hormonal profiles into a restricted range that becomes associated with specific behaviors and morphological traits.

Male mating success typically relies on secondary sexual characters that are paired with behaviors, such as territorial aggression and courtship displays. The hypothalamic–pituitary–gonadal (HPG) axis plays a pivotal role in the control of reproduction in vertebrates. At the apex of the HPG axis, gonadotropin-releasing hormone (GnRH) neurons secrete GnRH onto the pituitary in a pulsatile fashion. These pulses stimulate the rhythmic secretion of luteinizing hormone (LH) and follicle-stimulating hormone (FSH) from the pituitary into the blood (Fig. 1) [for details on birds see also (Li et al., 1994; Urbanski, 1984)]. The intermittent stimulation of gonadal tissue by gonadotropins then induces sex steroid synthesis and spermatogenesis in the male gonad.

**Fig. 1.**
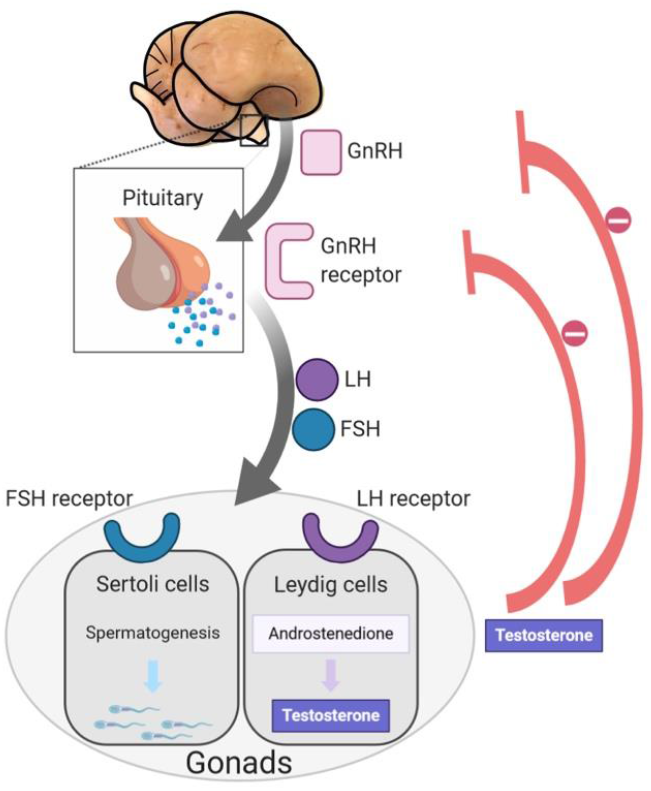
Overview of the hypothalamic-pituitary-gonadal (HPG) axis for testosterone synthesis. Gonadotropin-releasing hormone neurons in the anterior preoptic area of the Ruff brain send projections onto the pituitary where they release GnRH. The inset shows a close-up of the pituitary, which expresses a specific GnRH receptor (GNRH-R3). The binding of GnRH to this receptor triggers the production of the gonadotropins luteinizing hormone (LH) and follicle-stimulating hormone (FSH) that are released into the bloodstream. In the gonads, binding of FSH to its receptor in Sertoli cells stimulates spermatogenesis and binding of LH to its receptor in Leydig cells initiates a cascade of enzymatic steps that lead to the synthesis of androstenedione, which is then converted to testosterone. Testosterone begins its circulation through the bloodstream and serves to regulate GnRH, LH and FSH production through a negative feedback loop. (Created with BioRender).

Sex steroid hormones initiate and maintain social behaviors through genomic and non-genomic routes (Boonyaratanakornkit and Edwards, 2007). Among sex hormones, traditionally testosterone has been invoked to be largely responsible for differences in inter-male aggression (Goodson, 2005; Wingfield et al., 1987) although alternative more integrative mechanisms have been described recently (Goymann et al., 2019; Lipshutz et al., 2019; Wingfield et al., 2019).

Testosterone can reach the brain via the bloodstream or by *de-novo* synthesis in neural tissue (Baulieu, 1997; do Rego and Vaudry, 2016; Tsutsui et al., 1999). Yet, the regulation of social behavior does not only include testosterone but also its androgenic and estrogenic metabolites. The conversion of testosterone into estradiol is a requirement for mediating aggressive and sexual behavior in birds (Ball and Balthazart, 2004; Fusani et al., 2001; Steimer and Hutchison, 1980; Watson and Adkins-Regan, 1989) and estradiol is a known potent activator for sexual behavior in birds (Cornil et al., 2006). Estradiol can also be synthesized from estrone that is made from aromatized androstenedione, a testosterone precursor. Of note, androstenedione is only a weak activator of the androgen receptor, compared to testosterone. Another sex steroid, progesterone, has the ability to mediate aggression in rodent models [e.g. (Erpino, 1975; Erpino and Chappelle, 1971; Schneider et al., 2003; Yang et al., 2013)], male tree lizards [*Urosaurus ornatus* (Weiss and Moore, 2004)], as well as in females of the sex-role reversed black coucals (*Centropus grillii*) (Goymann et al., 2008). In the rufuos Hornero (*Furnarius rufus*) a South American ovenbird, the ratio between testosterone and progesterone has been shown to be a more sensible predictor for outcomes in short-term agonistic encounters than testosterone alone (Adreani et al., 2018).

The evolution of diversity in mating repertoires is largely facilitated by testosterone regulation (Ball and Balthazart, 2004; Balthazart et al., 2003; Day et al., 2007; Fusani et al., 2014; Oliveira et al., 2008). Therefore, knowledge of the physiological limits of androgen responsiveness is key to understand how behavioral diversity can evolve, especially in association with reproductive behaviors. Species that exhibit alternative reproductive tactics (ARTs) are ideal study systems for the proximate regulation of mating behavior because courtship, aggression and secondary sexual traits are dissociated from male gonadal function (Oliveira et al. 2008). GnRH challenges have proven particularly effective at unmasking when hormonal differences are largely regulated by the social context and interactions therein, and not by an actual physiological limitation to produce testosterone (Apfelbeck and Goymann, 2011; Barron et al., 2015; Cain and Pryke, 2017; Goymann and Flores Dávila, 2017; Goymann and Wingfield, 2004; Jawor et al., 2006; Spinney et al., 2006). During a GnRH challenge, a standardized amount of GnRH is administered to animals to examine the ability of the pituitary to respond to GnRH and the ability of the gonads to produce and secrete testosterone. In some species with reversible ranks or morphs, the entire HPG axis of subordinate males is suppressed by the social context. For example in the African cichlid *Astatotilapia burtoni*, the presence and agonistic behavior of dominant males actively keeps other males in a subordinate status (Maruska and Fernald, 2011). Subordinate males experience substantial shrinkage of GnRH neurons, decreased circulating 11-ketotestosterone and reduced spermatogenesis, which leads to smaller testes when the HPG axis is under social suppression for weeks (Davis and Fernald, 1990; Francis et al., 1993). However, this can be rapidly reversed if the social environment changes (Burmeister et al., 2005; Maruska et al., 2013). In contrast, in species with genetically determined ARTs, sneaker phenotypes are usually fixed for a lifetime (Oliveira et al., 2008). Such sneaker males tend to have low circulating levels of testosterone but their gonads are of similar size or even larger than those of territorial males, probably because this maximizes their fertilization success with fewer mating opportunities (Fu et al., 2001; Gross, 1996; Taborsky, 1998). This is possible because spermatogenesis actually requires low levels of testosterone to be initiated and maintained (Kustan et al., 2012; Oliveira et al., 2001, 2008). Understanding how genetic changes are able to produce fixed alternative morphs with distinct hormonal profiles is key to building the bigger picture of how morph-specific behaviors with a genetic basis can evolve.

A prominent inversion polymorphism that captures several genes involved in steroid synthesis and metabolism is responsible for three genetically determined male mating morphs - Independents, Satellites, Faeders - in the ruff, *Philomachus pugnax* (Küpper et al., 2016; Lamichhaney et al., 2016). During the breeding season, each morph has a distinct appearance and behavior (Hogan-Warburg, 1966; van Rhijn, 1973). Independent males have dark colored plumage and are overtly aggressive, whereas Satellites and Faeders, the two other ARTs, seldom exhibit aggressive behavior. Satellite males have light colored plumage and form temporary coalitions with Independent males. Faeders the smallest of the three morphs, lack ornamental plumage and do not perform male-typical courtship behaviors but rather adhere to a pure sneaking strategy (Jukema and Piersma, 2006). These morphological and behavioral differences are linked to striking hormonal differences in testosterone among the morphs: Independent males have high levels of testosterone compared to Satellites and Faeders (Küpper et al. 2016). Conversely, Satellites and Faeders have high levels of androstenedione, a testosterone precursor, compared to Independent males. Androstenedione, on its own, may act as a weak activator of androgen receptors and additionally, may be converted to testosterone or estrogen in target tissues including the brain (Soma et al., 2003). Indeed, experimentally elevating androstenedione in Independents results in increased aggression but is ineffective at eliciting any aggressive behavior in Satellites (albeit it increases courtship behaviour in Satellites, Morgan, 2009).

Remarkably, the three morphs are determined by a 4.5 Mb autosomal inversion. Independents have two copies of the ancestral non-inverted haplotype whereas Satellites and Faeders are both heterozygous for the inversion. Satellites and Faeders each have different inversion haplotypes: the Faeder inversion haplotype arose first and the Satellite haplotype is the result of subsequent recombination between the Faeder inversion and one or several ancestral Independent alleles (Küpper et al. 2016, Lamichhaney et al. 2016). The presence of several genes in the inversion region (i.e., *HSD17B2, SDR42E1, CY5B5*) with known involvement in sex steroid synthesis and metabolism suggests an inversion-based direct effect on hormone production and regulation. Yet, clear evidence of when and how the inversion alleles may disrupt sex steroid synthesis in inversion morphs remains elusive. Because the ability to produce testosterone may be a crucial trait directly controlled by genes within the inversion, identifying where within the HPG axis this disruption occurs is a necessary first step towards deciphering the underlying molecular mechanisms that generate morphological, neurobiological and behavioral differences across morphs.

In this study, we aimed to investigate the physiological and molecular mechanisms that maintain hormonal differences in adult male Ruffs during the breeding season. First, we performed a GnRH challenge to characterize the physiological range of testosterone that each morph can produce. We tested whether inversion morphs were capable of mounting a typical testosterone surge following a GnRH injection. We measured testosterone, androstenedione and corticosterone to investigate pituitary, gonadal and adrenal responses. Second, in a separate set of untreated birds, we measured the expression of key genes in the pituitary and gonads to assess hormonal sensitivity and production within the HPG axis. We measured the expression of genes required for pituitary sensitivity to sex steroids, gonadotropin-releasing hormone sensitivity (GnRH receptors), gonadotropin production (FSH) and gonadal sensitivity to gonadotropins (LH and FSH receptors), as well as the expression of the gene encoding the steroidogenic acute regulatory protein (StAR) which is essential for making cholesterol available for the synthesis of steroid hormones. We hypothesized that morph differences in the expression of any of these genes would indicate which tissue types and processes (i.e. hormone production or sensitivity) along the HPG axis would be responsible for the observed low circulating testosterone levels in inversion morphs.

## Materials and Methods

### Birds and housing

We used a total of 69 adult Ruff males (*Philomachus pugnax*) from a captive breeding flock at Simon Fraser University, with a mean age ± standard error (SE) of 5.44 ± 0.62 yrs. This captive population was originally established from eggs collected near Oulu, Finland, in 1985, 1989 and 1990 (Lank et al., 2013). We conducted all experiments and sample collection during three breeding seasons (see specific dates in relevant sections below). Males were housed in an outdoor aviary in a same-sex pen with visual access to females in adjacent pens prior to being tested and had unrestricted access to food and water at all times. Males devoted to tissue sampling were part of other experiments reported elsewhere. All housing and procedures (permit #1232B-17) were approved by the Animal Care Committee of Simon Fraser University operating under guidelines from the Canadian Council on Animal Care.

### GnRH challenge

We performed the GnRH challenge on 34 unique male birds (2018: 8 Independents, 2019: 11 Independents, 7 Satellites, 9 Faeders). In 2018, one Independent male with insufficient blood sampled was excluded for that year but re-tested in 2019. We performed these experiments on the following dates: in 2018 on June 11^th^ and 12^th^, and in 2019 on May 30^th^ and 31^st^, June 2^nd^ and June 11^th^. In 2018, we sought to test the efficacy of the dosage and the sampling time points, and we did not test all morphs. In both years, among Independents, we randomly assigned males to either the GnRH or the saline group. Because of sample size considerations and because we were interested in morph differences in the response to GnRH, we assigned all Satellites and Faeders to the GnRH group, similar to other studies (Cain and Pryke, 2017; Goymann and Wingfield, 2004). When possible, we daily injected the same proportions across treatment / morph groups to control for day effects. For each bird, the workflow was as follows. We captured the bird by net and brought it indoors, where we weighed and bled it from the left wing vein. The time from capture to first bleed was 4.17 ± 0.26 min (mean ± SE); this constituted the pre-challenge baseline sample. Immediately after blood sampling, we gave the bird an intramuscular GnRH1 injection (Bachem, H-3106) aimed at the pectoral muscle with a dose of 12.5 μg per 150 g body weight or a corresponding volume of saline, and placed it in a box that was kept indoors for the remainder of the experiment. We took two more blood samples at 30 and 90 min post-injection from the right wing and either right or left leg, respectively. We drew a maximum of 300 μl of blood each time, kept it on ice for the duration of testing time, and then centrifuged it at 3000 rpm for 15 min. We collected plasma and stored it at −20°C for up to four weeks before transporting it on dry ice to the lab in Germany and subsequent storage at −80°C until analysis.

### Hormone analysis

Plasma testosterone, androstenedione, progesterone and corticosterone concentrations were determined by direct radioimmunoassay (RIA) following previously published protocols (Goymann et al., 2001, 2006, 2008). For testosterone, androstenedione and corticosterone, plasma samples were extracted with dichloromethane (DCM) after overnight equilibration (4°C) of the plasma with 1500 dpm of tritiated testosterone, androstenedione, or corticosterone (Perkin Elmer, Rodgau, Germany). The organic phase was then separated from the aqueous phase by plunging the extraction tubes into a methanol-dry ice bath and decanting the dichloromethane phase into a new vial. This extraction step was repeated twice to increase extraction efficiency. Then, the DCM phase was dried under a stream of nitrogen at 40°C, dried samples resuspended in phosphate buffered saline with 1 % gelatine (PBSG) and left overnight at 4°C to equilibrate. Progesterone was extracted with ethyl ether after overnight equilibration (4°C) of the plasma with 1500 dpm of tritiated progesterone (Perkin Elmer, Rodgau, Ermany). Separation of the organic and aqueous phase was conducted similar to the procedure described for the other hormones. For all hormones, an aliquot of the redissolved samples was transferred to scintillation vials, mixed with 4 ml scintillation fluid (Packard Ultima Gold) and counted to an accuracy of 2-3 % in a Beckman LS 6000 β-counter to estimate individual extraction recoveries. The remainder was stored at −40°C until RIA was conducted. Mean ± sd extraction efficiency for plasma testosterone was 85.1 ± 3.6 % (N=157), for androstenedione 85.1 ± 3.9 % (N=120), for corticosterone 80.0 ± 9.6 % (N=80), and for progesterone 78.8 ± 4.9 % (N=42). For the RIA a standard curve was set up in duplicates by serial dilution of stock standard testosterone and androstenedione ranging from 0.39 – 200 pg, and corticosterone and progesterone from 1.95 – 1000 pg. Testosterone, androstenedione, corticosterone or progesterone antisera (all from Esoterix Endocrinology, Calabasas, CA, USA) were added to the respective standard curve, the controls and to duplicates of each sample (100μl). After 30 min the respective testosterone, androstenedione, corticosterone or progesterone labels were added and the assays incubated for 20 hours at 4°C. Then, bound and free fractions were separated at 4°C by adding 0.5 ml dextran-coated charcoal in PBSG assay buffer. After 14 min incubation with charcoal samples were spun (3600 g, 10 min, 4°C) and supernatants decanted into scintillation vials at 4°C. After adding 4 ml scintillation liquid (Packard Ultima Gold) vials were counted. Standard curves and sample concentrations were calculated with Immunofit 3.0 (Beckman Inc. Fullerton, CA), using a four parameter logistic curve fit. Data were analyzed in three assays for testosterone, two assays for androstenedione, and one assay for corticosterone and progesterone, respectively. The lower detection limits of the standard curves were determined as the first value outside the 95 % confidence intervals for the zero standard (Bmax) and was 3.6 – 4.9 pg/ml for testosterone, 6.7 – 8.1 pg/ml for androstenedione, 57.0 pg/ml for corticosterone, and 27.6 pg/ml for progesterone. The intra-assay coefficient of variation as determined by extracted chicken plasma pools was 1.5 %, 3.0 % and 10.4 % for the testosterone assays, twice 14.8 % for the androstenedione assays, 13.2 % for the corticosterone assay and 24.6 % for the progesterone assay. The inter-assay coefficient of variation as determined by extracted chicken plasma pools for the three testosterone assays was 5.9 % and for the two androstenedione assays was 1.3 %. Because the testosterone antibody used shows significant cross-reactions (44 %) with 5a-dihydrotestosterone our testosterone measurement may include a fraction of 5α-dihydrotestosterone.

### RNA extraction and cDNA synthesis

We extracted RNA from 35 pituitaries and 35 gonads (17 Independents, 9 Satellites, 9 Faeders) using the RNeasy Minikit (Qiagen) according to the manufacturer’s protocol. One modification for gonads was that 200 μl of the lysed homogenate was mixed with 300 μl lysis buffer, passed through a QIAShredder column to remove excess genomic DNA before starting the RNA extraction protocol. We measured RNA concentration with a Nanodrop and assessed RNA quality with the Bioanalyzer RNA nanochip (Agilent) in a subset of pituitary samples (N=25) and all gonad samples. We excluded two pituitary and four gonad samples from further analysis due to low yield or an RNA integrity number (RIN) below 5.4. For the remaining samples, the average RIN for pituitaries and gonads was 9.8 and 7.4, respectively. We synthesized a total of 500 ng (pituitary) and 1 μg (gonad) of RNA into cDNA using the iScript cDNA synthesis kit (Bio-Rad) in 20 μl reactions according to manufacturer’s instructions. We diluted cDNA ten-fold before use as template in qPCR assays.

### Primer design for target genes

To measure expression of genes essential to HPG axis function and steroid synthesis, we designed primers for ten genes. To assess pituitary sensitivity to GnRH we examined gene expression of gonadotropin-releasing hormone receptors (GNRHRs) I and III (hereinafter GNRH-R1 and GNRH-R3). Avian GNRH receptors have varying names in the literature [e.g. (Bédécarrats et al., 2006)]. Based on sequence identity conservation between ruff and chicken sequences we followed the nomenclature of (Joseph et al., 2009). We included the follicle-stimulating hormone beta subunit (FSHB) to gauge pituitary production of FSH and follicle-stimulating hormone receptor (FSHR) to assess gonadal sensitivity to FSH. Likewise, we designed primers for the gene encoding the luteinizing hormone receptor (LHR) to assess gonadal sensitivity to its corresponding gonadotropin. We selected the gene encoding the steroidogenic acute regulatory protein (StAR) because it could point to differences in cholesterol metabolism, which is essential for steroid synthesis. To test for sensitivity to sex steroids, we designed primers for genes encoding the estrogen receptors alpha and beta (ESR1 and ESR2), progesterone receptor (PGR) and androgen receptor (AR). For genes that we did not detect by name in the current ruff genome release (NCBI *Calidris pugnax* Annotation Release 100, software version 6.5) we searched for homologs using the respective chicken, turkey or quail sequences as query against the ruff genome with BLAST (Altschul et al., 1990). We designed primers with NCBI Primer-BLAST (Ye et al., 2012) and ensured that each amplicon spanned at least one exon-exon boundary and when possible we designed at least one primer within a given pair to anneal across an exon-exon junction. All primers and results from standard curves are listed in Table S1. Despite our best efforts, we were unable to locate the gene encoding the beta subunit of the luteinizing hormone with these methods, possibly because of the high GC content of this gene in avian species (chicken 72.12%, turkey 67.94%, quail 62.5%). Both FSH and LH use a common alpha subunit, but their respective beta subunits are hormone-specific (Proudman et al., 1999), therefore we could not use the alpha subunit as a proxy for LH production. We reasoned that even if we were able to bioinformatically retrieve this sequence in the Ruff, designing useful qPCR primers would have been difficult due to the numerous stretches of Gs and Cs.

### Primer design for reference genes

We tested a panel of 11 genes in gonad- and pituitary-derived cDNA for their suitability as reference genes for use in qPCR (**Table S1**, based on ((Loveland et al., 2014; St-Pierre et al., 2017; Zinzow-Kramer et al., 2014) and the best two genes for each tissue type that did not show differences between morphs were chosen after analysis with GeNorm and NormFinder.

### Quantitative PCR (qPCR)

We conducted and report all qPCR assays according to the Minimum Information for Publication of Quantitative Real-Time PCR Experiments (MIQE) guidelines (Bustin et al., 2009). We ran assays on a LightCycler 480 II (Roche) machine using the SsoAdvanced Universal SYBR Green Super mix (Bio-Rad) in 384-well plates (Roche) and each reaction was run in duplicate. We performed qPCR assays to generate the full data set across eight plates (each plate always balanced for morph) and standard curves on four plates. Each well consisted of a 10 μl reaction containing 1X SsoAdvanced Universal SYBR Green Super mix, 340 nM of each primer and either 3.75 ng (pituitary) or 7ng (gonad) of cDNA (i.e. 1.5 μl of the ten-fold cDNA dilution assuming a one-to-one correspondence between input RNA and synthesized cDNA). The cycling conditions were: pre-incubation step (95°C for 30 sec), 45 cycles (95°C for 10 sec, annealing and extension at 60°C for 30 sec) with acquisition at the end of each cycle, followed by a melt curve (95°C for 5 sec with 5 acquisitions per °C from 65°C to 97°C with a 0.11°C ramp rate). We performed calculations from the raw amplification data in the LightCycler 480 Software (version 1.5.1.62). On every plate we confirmed that each primer pair produced a single melt curve peak in the presence of cDNA template that was different from any primer dimer that may form when only water is used as template. Negative controls showed either no amplification or the presence of an unambiguous primer dimer peak. To test for any possible genomic carryover from RNA extraction to cDNA synthesis, we ran the qPCR with cDNA synthesis negative controls (i.e. no reverse transcriptase) for a subset of samples; as expected, none showed any amplification.

We ran a standard curve with serially diluted cDNA (1:10 to 1:100000) from one sample of an independent male to calculate the amplification efficiency of each primer pair, and efficiency was calculated separately for each tissue. This dilution range for *ESR2* primers was too wide, so for these primers we used serially diluted cDNA in a lower range (1:10 to 1:40). Efficiency was calculated with the Absolute Quantification tool and 2nd Derivative Maximum method which uses the formula [Efficiency = 10^-1/slope^] based on the quantification cycle (Cq, termed crossing point (Cp) in the software) and log concentration of template in each well. Efficiencies are listed in Table S1, all were above 1.92. We confirmed that the Cq values for the selected reference genes did not vary between Independents (n=5) and inversion morphs (n=6, 3 Satellites and 3 Faeders) (pituitary *HPRT1* Mann-Whitney p=0.6623, *RPL30* Mann-Whitney p=0.4286; gonads *HPRT1* t-test p=0.3797, *RPL32* t-test p=0.5002). We calculated the relative mRNA levels of target genes with the best reference genes for each tissue (*HPRT1* and *RPL30* for pituitaries, *HPRT1* and *RPL32* for gonads) with the Advanced Relative Quantification analysis, which is based on the ΔΔCT method by (Livak and Schmittgen, 2001) using the actual primer efficiencies from the standard curve instead of the preassigned value of 2.

### Statistical analyses

We used log-transformed hormone data for statistical analyses, but plotted non-transformed values in the figures. We calculated fold-change at the 30 min time point relative to baseline using non-transformed values. We used only Independent males to test for the effect of treatment (control vs. GnRH). We analyzed samples for testosterone in 2018 and 2019 and for androstenedione and corticosterone in 2019. In an initial model we used a two-way ANOVA on the difference between baseline and the 30 min time point to test for the main effect of year or interaction between year and treatment in testosterone levels. As neither the interaction between year and treatment nor year showed a significant effect (year x treatment F_(1,14)_=0.3959, p=0.593; year F_(1,14)_=1.921 p=0.1874; treatment F_(1,14)_=15.07, p=0.0017) we pooled the individuals by treatment group across both years for the final model.

To compare the hormonal changes between saline and GnRH-injected Independent males we prepared a mixed model with time, treatment and their interaction as fixed factors and ID as random factor. For posthoc tests, we tested whether the 30 and 90-min time points were different from baseline for each treatment group, and report Dunnett’s test statistics. To assess whether treatment groups differed clearly at each time point we used Sidak’s tests.

To assess hormonal variation across different morphs in response to the GnRH challenge, we compared GnRH-injected individuals only. We used a mixed model with morph, time and their interaction as fixed factors and ID as random factor. For posthoc tests, we tested whether the 30 and 90-min time points were different from baseline for each morph, and report Dunnett’s test. To test for morph differences at each time point we performed the Tukey’s test.

For each hormone, we analyzed morph differences as the relative change at 30 min from baseline (i.e. fold-change) with Kruskal-Wallis (KW) tests because sample sizes were too small to test for normality. We performed posthoc tests when the KW test was significant and used Dunn’s test to assess which morphs differed from one another.

For qPCR data, we analyzed variation in expression between morphs at each gene separately using either an ANOVA or Kruskal-Wallis test. For the analysis of sex steroid receptors in the pituitary, we used a Holm-Sidak test. All p values of posthoc tests are adjusted for multiple testing. We used GraphPad Prism (version 8) for statistical analyses.

## Results

### GnRH challenge in Independent males

The GnRH challenge resulted in different circulating testosterone concentrations in saline and GnRH-injected Independent males (Table S2, Fig. 2A). We found a strong interaction between time and treatment (F_(2,32)_=14.06, p<0.0001). Saline-injected birds had testosterone levels lower in post-injection samples relative to baseline, whereas, as predicted, GnRH-injected birds had higher levels of testosterone after 30-min post-injection relative to baseline (Dunnett’s p=0.0061) (Fig. 2A). We also found a treatment effect for circulating androstenedione concentrations (F_(1,9)_=8.129, p=0.019, Table S2, Fig. 2B). At the 30-min time point, GnRH-injected Independent males had higher androstenedione levels compared to baseline (Dunnett’s p=0.014), whereas saline-injected males did not differ between baseline and the 30-min sample (Dunnett’s p=0.971). At the 90-min time point there was no clear difference between treatment groups nor to the baseline measurements in androstenedione concentrations.

**Fig. 2.**
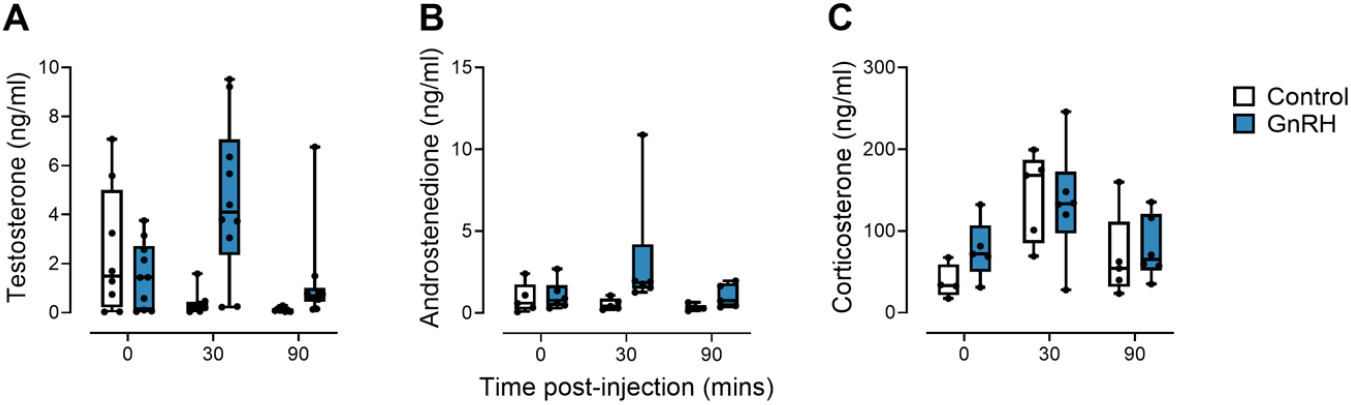
Hormone response in Independent Ruff males after a GnRH challenge. Time-course of steroid hormone levels in Independent males before and after a GnRH or control injection are shown for (A) testosterone, (B) androstenedione and (C) corticosterone after 0, 30 and 90 mins post-injection. Individual data points are superimposed on box plots. The “0” timepoint indicates pre-injection levels.

### Morph differences in the hormonal response to exogenous GnRH

#### Testosterone and androstenedione

Prior to injection, Independents had higher testosterone levels than Satellites and also higher, but not statically clearly, levels than Faeders (Table 1). Compared to Independent males, inversion morphs were incapable of mounting the typical robust testosterone increase after the GnRH challenge and at all sampling points inversion morphs showed a low testosterone level range (Fig. 3A, Table 1). Only Independent males presented a statistically clear increase in testosterone at 30-min after injection compared to their pre-injection levels (baseline vs. 30-min, Dunnett’s p=0.006, Table 1). At both post-injection time points inversion morphs had clearly lower levels than Independents (Table 1). We found clear differences in fold-changes for testosterone between morphs (Kruskal-Wallis, p=0.026). Mean fold-change (with range) relative to preinjection levels for inversion morphs were as follows: Satellites 1.89 (0.72–3.52), Faeders 1.38 (0.72–2.61), indicating a capacity to respond to GnRH, albeit to a lesser magnitude than Independents who mustered on average an 8.22 (0.995–45.62) fold-increase (Fig. 3B). The differences in fold-changes were clearest between Independents and Faeders (Dunnett’s p=0.022) but not statistically clear for comparisons involving Satellites (all p>0.05).

**Fig. 3.**
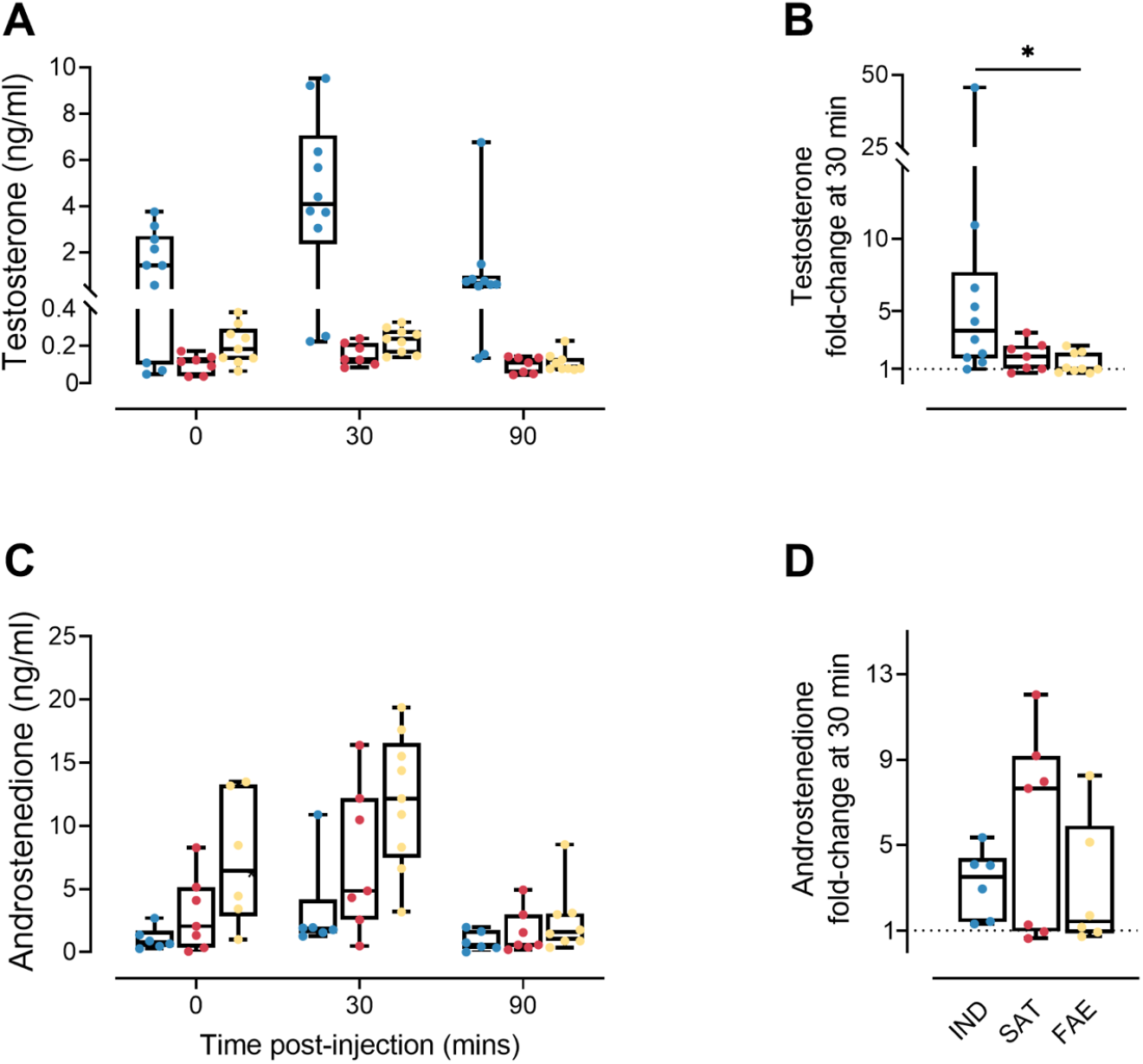
Ruff morphs differ in their androgenic response after a GnRH challenge. Hormone levels in Independent (blue), Satellite (red) and Faeder (yellow) males before and after a GnRH injection are shown for testosterone and androstenedione in top and bottom panels, respectively. Both Satellites and Faeders are heterozygous for a dominant autosomal inversion. Individual data points are superimposed on box plots. The time course from pre-to post injection is shown in (A and C) and fold-change at 30 min relative to pre-injection levels is shown in (B and D), the dashed line at ‘1’ fold-change indicates ‘no change’. The “0” time point indicates pre-injection levels. Box plots extend to the 25th and 75th percentiles, the line is drawn at the median and the whiskers extend to the minimum and maximum values. To provide informative detail to the illustration of testosterone levels for all morphs, note the introduction of y-axis breaks in A and B. See Table 1 for statistics on A, B. Asterisk denotes p<0.05.

**Table 1.**
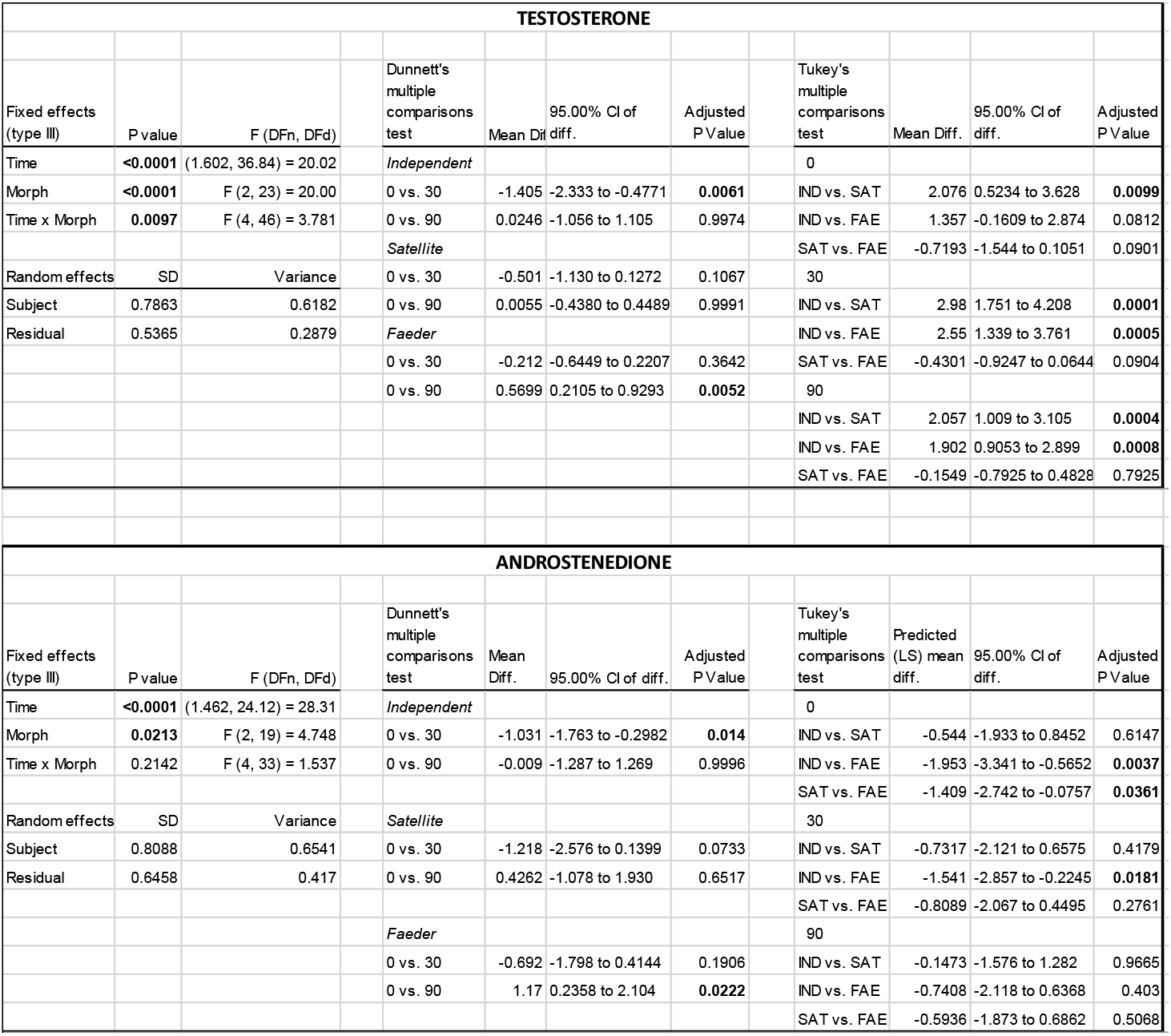
Morph differences in the hormonal response to exogenous GnRH: mixed effects model and posthoc test result

In contrast to the results for testosterone, inversion morphs at all sampling points had a higher average level of androstenedione compared to Independents (Table 1, Fig. 3C). The difference in androstenedione concentrations for Independents was statistically clear at 30 min compared to baseline (Table 1). However, the comparison of fold-change at 30 min relative to baseline revealed no clear differences among morphs (Kruskal-Wallis, p=0.619) (Fig. 3D), suggesting that the morph differences in androstenedione response are less pronounced than for testosterone.

### Corticosterone

Independents injected with GnRH did not differ in corticosterone levels from saline controls (treatment effect, F_(1,9)_=0.5035, p=0.496), but corticosterone levels increased with time (F_(1.638,13.10)_=19.77, p=0.0002) (Fig. 2C, Table S3). Post-injection levels at 30-min in both treatment groups were higher compared to their respective baseline but this was only statistically clear in the control group. We did not detect morph-specific corticosterone changes following the GnRH challenge (Table S4).

### Gene expression in pituitary and gonadal tissue

The three morphs did not clearly differ in expression of *GNRHR3*, *LHR*, *FSHR* or *FSHB* (Fig. 4, all tests p>0.05), key genes in the pituitary and gonads that regulate sensitivity to GnRH and gonadotropin sensitivity and production in our sample of untreated birds. In light of this, we hypothesized that differences in testosterone production were instead caused by functional differences in Leydig cells, in processes and pathways that occur after (i.e. downstream from) LHR activation (Fig. 5A). Consistent with this, morphs differed in *STAR* expression (ANOVA F_(2, 28)_=6.971, p=0.0035) (Fig. 5B). Yet, the observed *STAR* expression differences were not in the predicted direction. Faeders had the highest expression levels, exhibiting statistically clear differences compared to Independents (Tukey’s p=0.0024), but not to Satellites (Tukey’s p=0.087). There was no clear difference between Independents and Satellites (Tukey’s p=0.481). In agreement with previous reports, among the untreated birds (i.e. not part of the GnRH experiment) morphs differed in circulating testosterone levels (Kruskal-Wallis p=0.0049) (Fig. 5C): inversion morphs had similarly low levels (Satellite vs. Faeder, Dunn’s p>0.99) with Faeders (Dunn’s p=0.008), but not Satellites (Dunn’s p=0.064) exhibiting clearly lower levels than Independents.

**Fig. 4.**
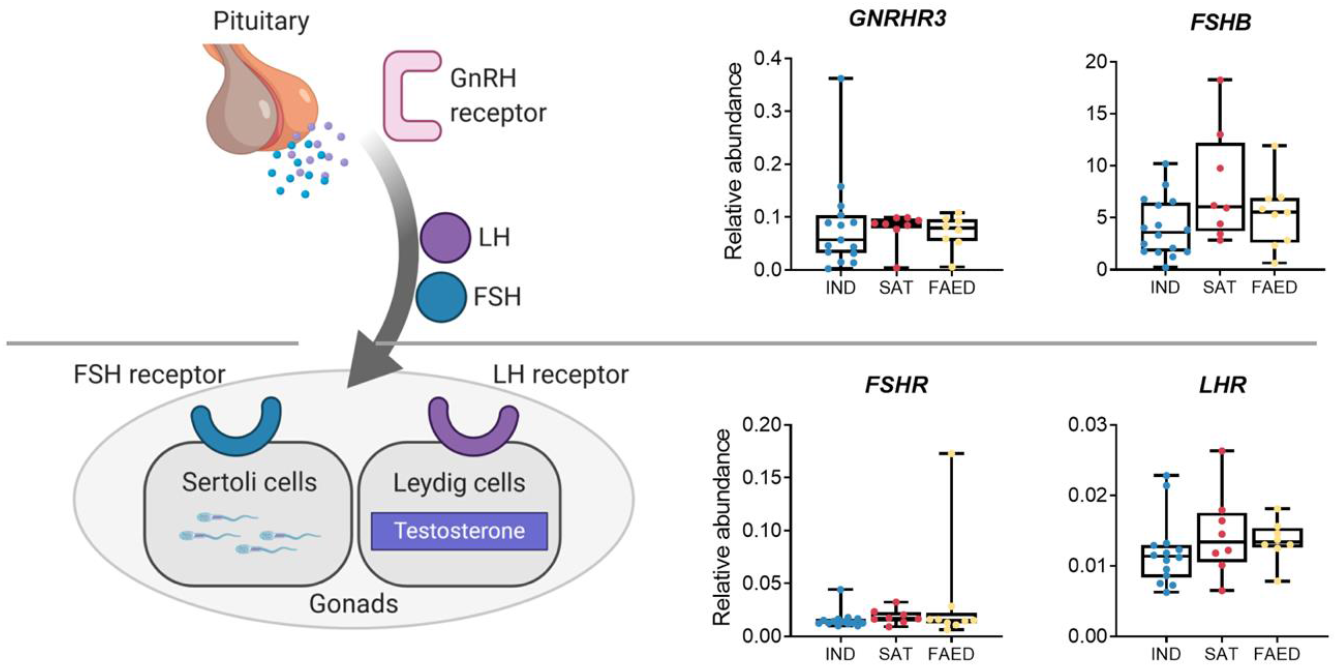
Ruff morphs do not differ in the expression of key genes in the pituitary and gonads that regulate gonadotropin sensitivity and production. Illustration of the end products for key genes responsible for pituitary sensitivity to GnRH and gonadotropin production and gonadal gonadotropin sensitivity (left panel). Box plots of *GNRHR3, FSHB, FSHR, LHR* expression in Independents (blue), Satellites (red) and Faeders (yellow) (right panel). Gene expression is plotted as abundance relative to two reference genes, see Methods for details. Box plots extend to the 25th and 75th percentiles, the line is drawn at the median and the whiskers extend to the minimum and maximum values. Individual data points are superimposed on box plots.

**Fig. 5.**
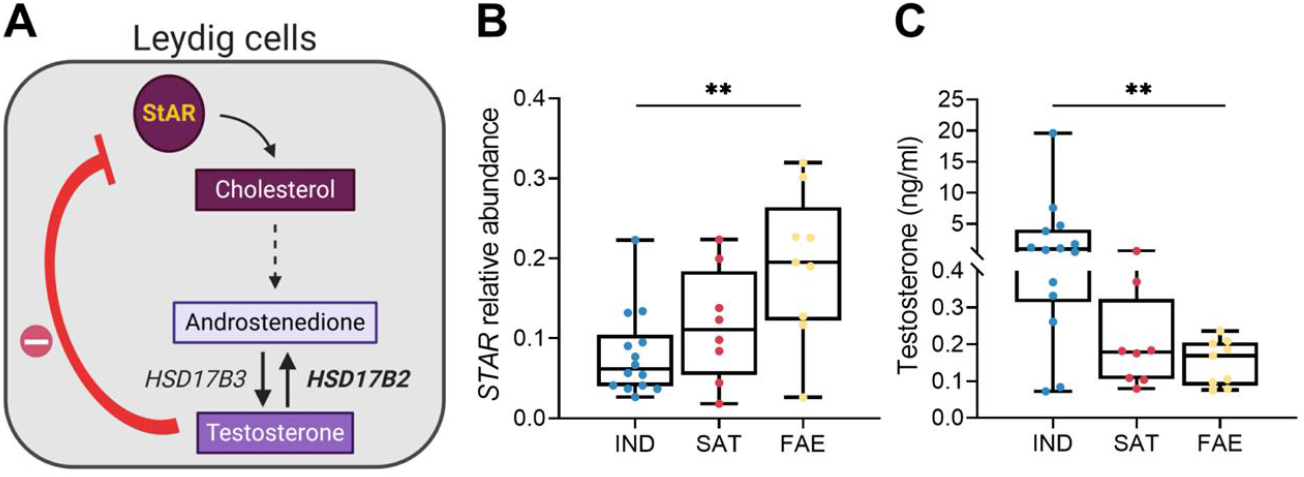
Gonadal STAR expression is atypical in Ruff inversion morphs during the breeding season. A) Schematic representation of regulatory interactions between StAR and testosterone in gonadal Leydig cells. A series of enzymatic reactions convert cholesterol into androstenedione denoted for simplicity by the dashed arrow. Androstenedione to testosterone conversion is reversible through HSD17B2 activity (bold), encoded by a gene inside the inversion shared by Satellites and Faeders. B) Morphs differed in levels of *STAR* expression C) in circulating testosterone. In both cases, only Faeders and Independents showed a clear statistical difference (asterisks, adjusted p < 0.01) from posthoc tests. According to feedback mechanisms between testosterone and *STAR* expression, inversion morphs, in particular Faeders, have such low levels of circulating testosterone that they are unable to down-regulate *STAR* expression in a typical manner. Box plots extend to the 25th and 75th percentiles, the line is drawn at the median and the whiskers extend to the minimum and maximum values. Individual data points are superimposed on box plots; panel C the y-axis break was introduced to provide informative detail on the full range of testosterone levels. (Created with BioRender).

Morphs differed in the expression of the progesterone receptor in the pituitary (ANOVA p-values Holm-Sidak adjusted: *PGR* p=0.0008; (Fig. 6A). Faeders had lower *PGR* expression than the other two morphs (Independents vs. Faeders, Tukey’s p=0.0001; Satellites vs. Faeders, Tukey’s p=0.0298). Satellites showed intermediate expression values, but the expression was not clearly different from that of Independents (Tukey’s p=0.281). None of the other of sex hormone receptors *ESR1*, *ESR2* or *AR* showed clear differences in pituitary expression among morphs (all p≥0.95). There was no relationship between *PGR* expression and circulating progesterone levels when all males were pooled (Spearman r=0.0046, p=0.98). However, within morphs we detected a positive relationship in Faeders (Spearman r=0.8095, p=0.0218), but not in the other two morphs (Satellites Spearman r=-0.028, p>0.99; Independents Spearman r=0.05, p=0.88) (Fig. 6B).

**Fig. 6.**
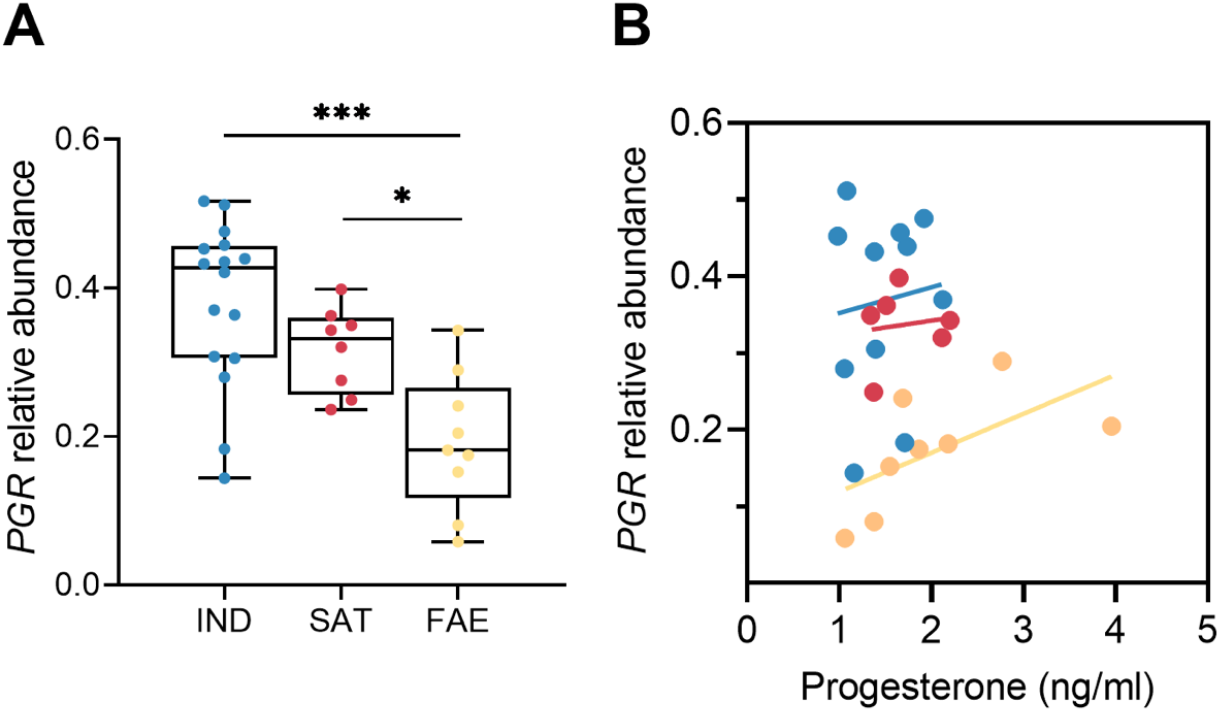
Morphs differ in progesterone receptor expression in the pituitary. A) *PGR* expression in Independents (blue), Satellites (red) and Faeders (yellow). Gene expression is plotted as abundance relative to two reference genes, see Methods for details. Box plots extend to the 25th and 75th percentiles, the line is drawn at the median and the whiskers extend to the minimum and maximum values. Individual data points are superimposed on box plots. B) Relationship between progesterone and PGR expression in each morph.

## Discussion

Breeding Ruffs show striking differences in behavior, appearance and androgen physiology across three morphs that only differ by a small autosomal inversion at the genomic level. Exactly how the inversion causes these hormonal and behavioral differences is unknown. In one set of birds, we tested the physiological range of androgenic production across the three morphs and in another set of untreated birds, we analyzed gene expression along the HPG axis to discern which tissue types and processes are likely to give rise to androgenic hormonal differences in this species. We found that the two inversion morphs, Satellites and Faeders, had generally a similar physiological response to the GnRH challenge and, unexpectedly, also showed similar expression profiles for most key genes within the HPG axis. This suggests that pituitary sensitivity to GnRH and gonad sensitivity to luteinizing hormone and follicle-stimulating hormone was similar across all morphs. In the GnRH challenge, both inversion morphs were incapable of synthesizing high levels of testosterone, but notably, were able to produce high levels of androstenedione.

In certain aspects our results parallel previous findings in the white-throated sparrow (*Zonotrichia albicollis*), a species with a two-morph system, ‘white’ and ‘tan’ striped, that are determined by the presence of a large inversion that contains approximately 1000 genes (Thomas et al., 2008; Thorneycroft, 1975). In this species, the white morph harbors the inversion and both males and females have brighter plumage, higher aggression and lower parental care during the breeding season compared to the tan morph, which is the presumed ancestral morph because it is homozygous for the non-inverted allele (Falls and Kopachena, 1993; Ficken et al., 1978; Horton et al., 2013). As in ruffs there are morph difference in testosterone levels: white morph males have higher testosterone levels than tan males. However, this difference is only seen in wild-caught birds, not in captive ones, perhaps because of differences in the stability of the social environment or induced stress, which could potentially suppress testosterone production via increased corticosterone (Spinney et al., 2006). In contrast, in Ruffs testosterone differences among morphs in the breeding season are highly pronounced even in captivity (Küpper et al. 2016, this study), suggesting that in the wild this difference may be even more pronounced and/or the effect of the inversion on circulating testosterone levels is less labile to the influence of the social environment. In white-throated sparrows, white-striped morph males also show a more robust increase in testosterone after a GnRH challenge compared to tan-striped males even though the LH surge caused by the injection is not different between the morphs (Spinney et al., 2006). In that study, the authors suggested that testosterone differences might stem from morph differences in the expression of LHR and/or steroidogenic enzymes in the gonads. In contrast, we confirmed in ruffs that there are no morph differences in gonadal *LHR* expression despite the presence of a subdued testosterone response of the inversion morphs compared to that of Independents. To what extent steroidogenic enzymatic activity in ruffs is affected by the inversion is still not known.

We predicted that if the cause of low testosterone in inversion morphs were due to a direct constraint from the inversion itself, testosterone levels after GnRH administration would show either a subdued increase or no increase at all in Satellites and Faeders. Alternatively, if low testosterone in inversion morphs were regulated by the social context, we expected that testosterone levels in inversion morphs would increase after the GnRH injection. Our results provide strong support for the former, that is, a direct effect of the inversion on the capacity to produce testosterone. This suggests that the inversion irreversibly disrupts testosterone synthesis in Faeders and Satellites. Our results from the expression of key genes along the HPG axis in untreated males point towards the gonads as the most likely site of physiological differentiation that leads to the observed low circulating testosterone levels in inversion morphs. First, we found that expression in the pituitary is broadly consistent with no differentiation in pituitary function between morphs, though Satellites did show a trend for higher *FSHB* expression. However, we cannot rule out the possibility that there are morph differences in the amount of GnRH that is naturally produced, as we have not studied properties of GnRH neurons or endogenous GnRH production. In other species with fixed ARTs, differences in the number and size of GnRH neurons between morphs exist suggesting that there might also be differences in how much GnRH is synthesized and released. For example, in the white-throated sparrow, females of the inverted morph have fewer, but larger, GnRH-immunoreactive neurons in the septo-preoptic area (Lake et al., 2008). Likewise, in the plainfin midshipman, *Porichthys notatus*, territorial courting males have larger GnRH neurons than type II non-territorial sneakers (Grober et al., 1994). However, we did not find differences in *GNRH-R3* expression between morphs, which would have been indicative of morph differences in GnRH receptor expression and/or endogenous GnRH production.

In birds, two GnRH isoforms (*GnRH1* and *GnRH2*) and two genes for GnRH receptors exist (*GNRH-R1* and *GNRH-R3*). Originally, we had intended to measure expression of both GnRH receptors to determine whether both were expressed in the pituitary and whether one was more abundant than the other. In chicken (*Gallus gallus*), the GnRH-R3 receptor is thought to be the major receptor for regulating pituitary gonadotroph function, in part because its levels in the pituitary are 1400-fold higher than GnRH-R1 (Joseph et al., 2009). The primers we designed for *GNRH-R1* did not pass quality control criteria, most likely because this gene has low expression (see Table S1). Therefore, we were unable to examine *GNRH-R1* expression in the pituitary. However, in line with the study by Joseph et al. (2009) we observed *GNRH-R3* expression levels similar to the expression seen in chicken, suggesting that *GNRH-R3* is the major functional receptor for GnRH1 in the pituitary.

Second, we found that, among untreated males, both inversion morphs had an apparent chronic upregulation of *STAR* expression in the gonads. Genes early in a biosynthetic pathway can be major regulators of the synthesis of particular end products, in part due to negative feedback mechanisms that ensure prompt and tight regulation (Manna and Stocco, 2005). StAR is essential to making cholesterol available for the synthesis of sex steroids (Lin et al., 1995) (Fig. 5A). To achieve this, the StAR protein transports cholesterol from the outer to inner mitochondrial membrane and then from that point on, through a series of enzymatic steps cholesterol can be converted into any of several steroid hormones (Stocco, 2001). In vertebrates, the *STAR* gene is typically regulated by several hormones, including testosterone, and its function has been suggested to be a rate-limiting step for testosterone synthesis, despite not being a ‘strict’ enzymatic protein (Houk et al., 2004; Manna and Stocco, 2005). In the dark-eyed junco *(Junco hyemalis) STAR* expression levels predict individual variation in testosterone in males, with higher expression in males with relatively high circulating testosterone levels (Rosvall et al., 2016). Accordingly, we had initially hypothesized that inversion morphs would have reduced *STAR* expression compared to Independent males. However, the relatively high levels of androstenedione in inversion morphs added complexity to this prediction. Our *STAR* expression result suggests that inversion morphs have the substrate needed (i.e. cholesterol) to mount a steroidal response (also evidenced by GnRH-treated inversion morphs that were able to elevate androstenedione levels). Yet still, elevated *STAR* levels even in untreated inversion morphs is indicative of a failed attempt to elevate testosterone levels.

One reason for the low circulating levels of testosterone could be an increased back conversion from testosterone into androstenedione in both inversion morphs (Fig. 5A). Inversion morphs have high levels of androstenedione during the breeding season (Küpper et al. 2016, this study). Intriguingly, the gene encoding *HSD17B2*, an enzyme that converts testosterone to androstenedione is located within the inversion region and its immediate surrounding sequence contains several deletions in the inversion haplotypes (Küpper et al., 2016; Lamichhaney et al., 2016). Interestingly, we observed a moderate increase of androstenedione levels in both inversion morphs 30 mins post-injection of GnRH. This could indicate that in the inversion morphs testosterone is converted back to androstenedione rather quickly, such that at our sampled time points the short-lived transient increase in testosterone in inversion morphs had passed already. However, we think that such a scenario is unlikely as gonadal *HSD17B2* levels are similar across morphs (Loveland et al., in preparation). Rather we suggest that the lack of a robust increase in testosterone in inversion morphs after a GnRH injection results from an existing impairment in androstenedione to testosterone conversion leading to a lack of testosterone availability in the first place.

We report a novel link between progesterone receptor expression in the pituitary and circulating progesterone in the Faeder morph, which has the most ancient inversion. This result while somewhat unexpected, offers an interesting avenue for future exploration. *PGR* transcript abundance was highest in Independents followed by Satellites and then Faeders; and only in Faeders, *PGR* transcript abundance was positively correlated to circulating progesterone levels. Assuming the same binding properties between progesterone and its receptor among morphs, a reduction in PGR would require a greater amount of progesterone in Faeders for the same receptor occupancy [i.e. “law of mass action” (Nussey and Whitehead, 2001)]. In black coucals, a sex-role reversed species, progesterone inhibits aggression in females (Goymann et al., 2008) and in the rufous Hornero, the ratio between testosterone and progesterone can predict outcomes in shortterm agonistic encounters better than testosterone alone (Adreani et al., 2018). In tree lizards, fixed color morphs result from organizational effects of both progesterone and testosterone levels and most striking, is that experimental manipulation of progesterone levels with a single injection on the day of hatching is sufficient to induce males to become the orange-blue morph (Moore et al., 1998). In another study, progesterone implants in castrated males triggered aggressive behavior, albeit to a lesser degree than testosterone implants (Weiss and Moore, 2004). While this evidence implicates progesterone in aggression regulation in lizards and birds, whether and how pituitary PGR expression and Faeder behavior and physiology could be linked is less clear. Perhaps, because Faeder males have been subjected to selection that favors a non-aggressive phenotype, the progesterone receptor may have lost any functional role in relation to male aggression altogether in this morph and as such, its expression is reduced.

Given the inversion is the main location of genetic differentiation among morphs, we expect that constraints on testosterone production are life-long. Accordingly, inversion morph males will already experience significant differences in exposure to androgens early in development, compared to Independent males. In birds, how the hormonal environment in early development affects the sexual differentiation of the brain and behavior is complex and species-specific (Adkins-Regan, 1987; Balthazart and Ball, 1995; Gahr, 2004). To date, this has not been investigated in Ruffs. Nonetheless, we expect brain development to be influenced by an interaction between the hormonal environment, sex chromosomes and inversion genotype, ultimately leading to adult neural circuits for morph-specific behaviors. The earliest detected differences between Ruff morphs are between chicks. At hatching, chicks of all morphs are similar in size but quickly develop different growth trajectories (Giraldo-Deck et al., 2020). Still, behavioral differences across male morphs may be influenced by organizational effects of testosterone. Consistent with a hypothesis for strong organizational effects, wherein de-masculinization of the brain occurs due to estrogen levels in females, adult Ruff females that are given testosterone implants grow male-like plumage and exhibit male-typical behaviors according to their genotype (Lank et al., 1999). These results demonstrate that genotype-specific neural circuits for male behaviors are present in both sexes and can be activated by testosterone. In addition, this suggests that since the inversion is present in all cells, the same constraints that lead to low circulating testosterone levels in inversion morphs could also be present in the *de-novo* production of testosterone in neural tissue, which could further accentuate neurobiological differences among morphs.

Sneaker males may have evolved to have high levels of glucocorticoids as an adaptive response that enables the maintenance of a stable mating phenotype (Arterbery et al., 2010; Knapp and Neff, 2007). Furthermore, it is possible that acute increases in androgens are also accompanied by increased corticosterone. We therefore investigated whether morphs differ in the rate at which they can increase and decrease levels of this stress hormone. However, we did not find morph-specific differences, rather corticosterone increased in both saline and GnRH injected groups. This suggests that Ruff morphs experienced handling stress in a similar way.

In the white-throated sparrow, recent studies have elucidated in great detail how two genes from within the inversion, estrogen receptor alpha (*ESR1*) and vasoactive intestinal peptide (*VIP*), are responsible for morph behavioral differences in aggression, song frequency and parental care in specific regions of the brain (Horton et al., 2020, 2014a; Merritt et al., 2020; Sun et al., 2018). Notably, because breeding season testosterone differences between morphs in the white-throated sparrow disappear under captive social settings, the picture that has emerged is that these key genes within the inversion affect neural gene expression that is associated with estrogen sensitivity and production of the VIP hormone is specific brain areas. But importantly, sex steroid hormone synthesis is not as strongly influenced by the inversion than as we report here for the Ruff. Thus, from these two exemplar fixed ART species we can appreciate how inversions, and the genes within them, can evolve to have variation in expression due to sequence divergence in regulatory regions that are the consequence of suppressed recombination (e.g. *cis*-effects in *ESR1* and *VIP* in *Z. albicollis*) as well as trans-effects, where the presence of the inversion affects the expression of genes that are outside of the inversion and functionally associated with androgen production.

## Conclusion

We investigated how inversion morphs respond hormonally to exogenous GnRH in male Ruffs. We found that the ability to synthesize testosterone is severely disrupted in the two inversion morphs, Faeders and Satellites. There was no evidence for widespread reduced functioning or throughput in early stages of the testosterone synthesis pathway, as inversion morphs show over-expression of *STAR*, which provides substrate for synthesis of sex hormones. Given the absence of clear differences across morphs in the expression of key genes in the pituitary for HPG axis function (see Fig. 4) we argue that morph differences in the capacity to produce testosterone originate within the gonads, and notably, are not due to morph differences in gonadotropin production nor sensitivity. In support, we found that inversion morphs were able to produce a transient increase in androstenedione, a testosterone precursor, following the GnRH injection supporting the view that their sensitivity to GnRH is comparable to that in Independents. Taken together, our study shows that the chromosomal inversion may have profound effects on the expression of genes involved in steroid regulation that are located outside of the inversion region.

## Acknowledgements

We would like to thank Stephanie Roilo for bird care and technical assistance in sample collection, Aaron Walchuk, Soma Marton, Gabi Trainor for aviary maintenance, Soma Marton for technical assistance in the GnRH challenge experiment, Hubert Schwabl for discussions in the early planning stages of this project, Monika Trappschuh for hormone sample processing and Antje Bakker for assistance with the Bioanalyzer.

## Author contributions

JLL conceived the GnRH-related and gene expression elements of the project; JLL, CK and DL designed experiment conditions and sample collection; JLL, LG and DL performed the GnRH challenge experiment; WG supervised hormone measurements; MG provided lab space and reagents, JLL collected and processed all samples and analyzed the data. JLL wrote the initial draft with input from CK, all authors edited the manuscript.

