## Supplemental Table 1 for "Functional differences in the hypothalamic-pituitary-gonadal axis are associated with alternative reproductive tactics based on an inversion polymorphism"

| Gene | Accession | Primer name | Primer sequence | Exons spanned | Standard Curve | | | | | | | | Other notes |
|---|---|---|---|---|---|---|---|---|---|---|---|---|---|
| | | | | | Gonad | | | | Pituitary | | | | |
| | | | | | Efficiency | Slope | Error | Y-intercept | Efficiency | Slope | Error | Y-intercept | |
| *GAPDH* | XM_014957084.1 | gapdhFOR1<br>gapdhREV1 | TCCCATGTTCGTCATGGGTG<br>GATGGCATGGACAGTGGTCA | 4-6 | 1.959 | -3.424 | 0.003 | 17.330 | 1.935 | -3.489 | 0.0080 | 15.71 | |
| *RPL30* | XM_014965539.1 | rpl30FOR1<br>rpl30REV1 | CTCTAGGCTTCAGCTGGTTATG<br>TCGATCTCCGATTTCCTCAAAG | 3,4 | 1.936 | -3.485 | 0.004 | 16.690 | 1.976 | -3.381 | 0.0047 | 17.04 | |
| *RPS13* | XM_014962266.1 | rps13FOR1<br>rps13REV1 | CCACGTGGCTGAAACTTACT<br>CCCTCAGGATCACACCAATTT | 2-4 | 1.971 | -3.393 | 0.000 | 17.720 | 1.983 | -3.362 | 0.0018 | 17.86 | |
| *18S* | XM_014962545.1 | 18S_F1201<br>18S_R1321 | GGCCCGAAGCGTTTACTTTG<br>CCCCGTTTCCGAAAACCAAC | NA | 1.986 | -3.355 | 0.004 | 4.496 | 1.977 | -3.379 | 0.0074 | 4.686 | |
| *HMBS* | XM_014949590.1 | hmbs_F2206<br>hmbs_R2329 | AGACCGGCGTTCATTATCCC<br>TCAACCCCAAGGTTCTCTGC | 11,12 | 1.920 | -3.513 | 0.020 | 19.840 | 1.978 | -3.376 | 0.081 | 22.800 | |
| *HPRT1* | XM_014963124.1 | hprt1_F522<br>hprt1_R602 | AAGAACTCCTCGAAGCGTGG<br>AGGGCGTATCCCACAACAAA | 7,8 | 1.985 | -3.358 | 0.008 | 20.990 | 2.128 | -3.049 | 0.0092 | 22 | |
| *RPL4* | XM_014946127.1 | rpl4_F502<br>rpl4_R582 | AACTCAGAAGCGTTACGCCA<br>ATGCGGTGACCTTTGGACAT | 4,5 | 1.962 | -3.417 | 0.011 | 19.150 | 2.014 | -3.29 | 0.0156 | 18.46 | |
| *RPL32* | XM_014946777.1 | rpl32_F192<br>rpl32_R266 | ACCAAGAAGTTCATCCGCCA<br>GATACCCCTCGGTTTACGCC | 2,3 | 2.009 | -3.300 | 0.002 | 17.510 | 1.92 | -3.53 | 0.0171 | 16.77 | |
| *ACTIN* | XM_014936234.1 | actb_F27<br>actb_R157 | GATGATGATATCGCCGCGCTC<br>ACCATCACACCCTGATGTCTG | 1,2 | 1.973 | -3.389 | 0.003 | 18.720 | 1.959 | -3.425 | 0.0070 | 18.48 | |
| *YWHAZ* | XM_014965334.1 | ywhaz_F215<br>ywhaz_R346 | ACGGAAGGCGCTGAGAAAAA<br>TGCTTGTGAAGCATTGGGGA | 1,2 | 1.973 | -3.389 | 0.011 | 18.430 | 2.021 | -3.273 | 0.0267 | 19.59 | |
| *PPIH* | XM_014946915.1 | ppih_F65<br>ppih_R156 | CCCAACAACCCTGTGGTCTT<br>TTGGGTACAACGTCAGCGAA | 1,2 | 1.980 | -3.371 | 0.022 | 20.830 | 1.955 | -3.434 | 0.0235 | 21.58 | |
| *FSHB* | XM_014949257.1 | FSHB_F148<br>FSHB_R344 | TGCTTCACAAGGGATCCAGTA<br>CTCACAGTGCAGTCAGTGGT | 1,2 | NA | NA | NA | NA | 1.953 | -3.441 | 0.002 | 17.720 | |
| *GNRH-RI* | XM_014946100.1 | GNRHRI_F745<br>GNRHRI_R825 | GCTTCACCCCAGCTTTTCCTT<br>GAAGCTCCCATGAGTAACGC | 2,3 | NA | NA | NA | NA | | | | | Inconsistent amplification of duplicate reactions. Most wells show two peaks, often with same height so unclear if one is primer dimer. Literature suggests this R1 is not highly expressed in |
| *GNRH-RI* | XM_014946100.1 | GNRHRI_F746<br>GNRHRI_R851 | CTTCACCCCAGCTTTTCCTTTT<br>TAGACAGTTTCCTCCCAGTGC | 2,3 | NA | NA | NA | NA | | | | | Inconsistent amplification of duplicate reactions. Some wells show single peak, some show double peak, too inconsistent |
| *GNRH-RIII* | XM_014952694.1 | GNRHRIII_F891<br>GNRHRIII_R1037 | ATCATGGTCTGCTGCTACGC<br>GACCAGGCTCATCTTCAGCA | 3,4 | NA | NA | NA | NA | 1.974 | -3.386 | 0.036 | 24.190 | |
| *GNRH-RIII* | XM_014952694.1 | GNRHRIII_F689<br>GNRHRIII_R808 | GAGGAACCGCATCATGCTCTA<br>GTGGTGCACTGGGTGAAGTT | 2,3 | NA | NA | NA | NA | 2.014 | -3.288 | 0.073 | 23.780 | |
| *PGR* | XM_014965714.1 | PGRFOR1<br>PGRREV1 | AAGGGGTTGTGGCCAATTCA<br>TCTCGGGGAATTCAACGCTC | 7,8 | 2.103 | -3.097 | 0.021 | 26.000 | 2.008 | -3.302 | 0.0021 | 20.86 | |
| *ESR2*[a] | XM_014951586.1 | ERbetaFOR1<br>ERbetaREV1 | GGGGTCTGGTCATGTGAAGG<br>CATTTCCGTAGCCGACATGC | 3,4 | 2.074 | -3.156 | 0.0423 | 25.59 | | | | | |
| *ESR1* | XM_014951246.1 | ERalphaFOR1<br>ERalphaREV1 | GTCTGGTCTTGTGAGGGCTG<br>TTCTTAGTCGGCAAGCCTGG | 2,3 | 2.001 | -3.319 | 0.060 | 25.660 | 1.997 | -3.328 | 0.0420 | 20.43 | |
| *AR* | XM_014954960.1 | ARFOR1<br>ARREV1 | GTTCGGGTGGCTTCAGATCA<br>GCTTCTGGTTCTTCAGGCCA | 6,7 | 1.954 | -3.437 | 0.011 | 24.640 | 1.948 | -3.454 | 0.0170 | 21.64 | |
| *FSHR* | XM_014962363.1 | FSHR_F458<br>FSHR_R551 | CACAAGGTGCACTCTCTCCA<br>AACTCAGGCCCGTGAATGAA | 5,6 | 2.107 | -3.090 | 0.002 | 22.980 | NA | NA | NA | NA | |
| *LHR* | XM_014962370.1 | LHR_F502<br>LHR_R571 | AACAATGAATCGCTGACACTCAA<br>CATTGAAGGCATGGCTGTGG | 6,7 | 2.012 | -3.292 | 0.019 | 25.700 | NA | NA | NA | NA | |
| *STAR* | XM_014958429.1 | staRqF1<br>staRqR1 | ATCAGGGCAGAGAATGGTCC<br>TGATGATGGTCTTCGGGAGC | 5-7 | 1.951 | -3.445 | 0.011 | 23.270 | NA | NA | NA | NA | |

a = The primer efficiency calculated from gonad RNA was used for analysis of pituitary data i.e. a separate primer efficiency calculation for pituitary was not performed

NA = not applicable
