## Supplemental Tables 2-4 for "Functional differences in the hypothalamic-pituitary-gonadal axis are associated with alternative reproductive tactics based on an inversion polymorphism"

**Table S2. Effects of GnRH on androgens in Independents: Mixed effect model and posthoc test results (related to Fig. 2)**

| TESTOSTERONE | | | | | | | | | | | | |
|---|---|---|---|---|---|---|---|---|---|---|---|---|
| Fixed effects (type III) | P value | F (DFn, DFd) | | Dunnett's multiple comparisons test | Mean Diff. | 95% CI of diff. | Adjusted P Value | | Sidak's multiple comparisons test | Mean Diff. | 95% CI of diff. | Adjusted P Value |
| Time | **0.0011** | F (1.314, 21.02) = 12.09 | | *Control* | | | | | | | | |
| Treatment | **0.0293** | F (1, 16) = 5.731 | | 0 vs. 30 | 1.268 | 0.1855 to 2.350 | **0.026** | | Control - GnRH | | | |
| Time x Treatment | **<0.0001** | F (2, 32) = 14.06 | | 0 vs. 90 | 2.182 | 0.6274 to 3.737 | **0.0112** | | 0 | 0.1663 | -2.362 to 2.695 | 0.9973 |
| | | | | *GnRH* | | | | | 30 | -2.507 | -4.108 to -0.9056 | **0.0022** |
| Random effects | SD | Variance | | 0 vs. 30 | -1.41 | -2.333 to -0.477 | **0.0061** | | 90 | -1.991 | -3.120 to -0.8622 | **0.0008** |
| Subject | 1.185 | 1.405 | | 0 vs. 90 | 0.025 | -1.056 to 1.105 | 0.9974 | | | | | |
| Residual | 0.7974 | 0.6358 | | | | | | | | | | |

| ANDROSTENEDIONE | | | | | | | | | | | | |
|---|---|---|---|---|---|---|---|---|---|---|---|---|
| Fixed effects (type III) | P value | F (DFn, DFd) | | Dunnett's multiple comparisons test | Mean Diff. | 95% CI of diff. | Adjusted P Value | | Sidak's multiple comparisons test | Mean Diff. | 95% CI of diff. | Adjusted P Value |
| Time | 0.1087 | F (1.831, 15.56) = 2.610 | | *Control* | | | | | | | | |
| Treatment | **0.0191** | F (1, 9) = 8.129 | | 0 vs. 30 | 0.145 | -2.255 to 2.544 | 0.9714 | | Control - GnRH | | | |
| Time x Treatment | 0.206 | F (2, 17) = 1.736 | | 0 vs. 90 | 0.567 | -1.055 to 2.189 | 0.4725 | | 0 | -0.4601 | -2.642 to 1.722 | 0.8899 |
| | | | | *GnRH* | | | | | 30 | -1.636 | -2.916 to -0.3547 | **0.014** |
| Random effects | SD | Variance | | 0 vs. 30 | -1.03 | -1.763 to -0.298 | **0.014** | | 90 | -1.036 | -2.347 to 0.2749 | 0.1261 |
| Subject | 0.4164 | 0.1734 | | 0 vs. 90 | -0.01 | -1.287 to 1.269 | 0.9996 | | | | | |
| Residual | 0.7369 | 0.5431 | | | | | | | | | | |

### Table S3. Test for effects of GnRH on corticosterone in Independents

| **CORTICOSTERONE** | | | | **CORTICOSTERONE** | | | |
|---|---|---|---|---|---|---|---|
| Fixed effects (type III) | P value | F (DFn, DFd) | | Dunnett's multiple comparisons test | Mean Diff. | 95% CI of diff. | Adjusted P Value |
| Time | **0.0002** | F (1.638, 13.10) = 19.77 | | *Control* | | | |
| Treatment | 0.4959 | F (1, 9) = 0.5035 | | 0 vs. 30 | -1.37 | -2.018 to -0.718[illegible] | **0.0064** |
| Time x Treatment | 0.0665 | F (2, 16) = 3.227 | | 0 vs. 90 | -0.49 | -1.207 to 0.226[illegible] | 0.1239 |
| | | | | *GnRH* | | | |
| Random effects | SD | Variance | | 0 vs. 30 | -0.49 | -1.235 to 0.257[illegible] | 0.1569 |
| Subject | 0.4994 | 0.2494 | | 0 vs. 90 | -0.02 | -0.8200 to 0.77[illegible] | 0.9928 |
| Residual | 0.3188 | 0.1017 | | | | | |

#### Table S4. Test for morph differences in corticosterone response after GnRH challenge

| CORTICOSTERONE | | | | | | | |
|---|---|---|---|---|---|---|---|
| Fixed effects (type III) | P value | F (DFn, DFd) | | Dunnett's multiple comparisons test | Mean Diff. | 95.00% CI of diff. | Adjusted P Value |
| Time | **0.006** | F (1.940, 29.10) = 6.213 | | *Independent* | | | |
| Morph | 0.5083 | F (2, 19) = 0.7014 | | 0 vs. 30 | -0.4888 | -1.235 to 0.2577 | 0.1569 |
| Time x Morph | 0.1954 | F (4, 30) = 1.618 | | 0 vs. 90 | -0.0236 | -0.8200 to 0.7728 | 0.9928 |
| | | | | *Satellite* | | | |
| Random effects | SD | Variance | | 0 vs. 30 | -0.9591 | -1.882 to -0.03617 | **0.0438** |
| Subject | 0.2537 | 0.06436 | | 0 vs. 90 | -0.6551 | -1.781 to 0.4713 | 0.2046 |
| Residual | 0.4696 | 0.2206 | | *Faeder* | | | |
| | | | | 0 vs. 30 | -0.2132 | -1.170 to 0.7437 | 0.7371 |
| | | | | 0 vs. 90 | -0.2441 | -0.8504 to 0.3623 | 0.4285 |
